# Circadian Variation of Salivary Oxytocin in young adult women

**DOI:** 10.1101/2025.03.26.645423

**Authors:** M. Teixeira de Almeida, L. Quattrocchi, N. Perroud, T. Aboulafia-Brakha

**Affiliations:** Faculty of Medicine, University of Geneva; Department of Psychiatry, Geneva University Hospitals

**Keywords:** Circadian variation, oxytocin, salivary biomarkers, borderline personality disorder, hormonal cycle, female participants, young adult women

## Abstract

This article presents circadian variation in salivary oxytocin levels in a sample of 91 female participants, including 47 healthy controls and 44 patients with borderline personality disorder (BPD). A significant increase in salivary oxytocin levels was observed between awakening and early afternoon. There were no significant group differences and no Group × Time interaction. These findings have implications for research conducted in the field and suggest the need to control for time of assessment, as done in studies assessing cortisol.

## Introduction

The neuropeptide oxytocin is increasingly being studied for its role in social bonding, emotion regulation, stress response, and cognitive flexibility [1-5]. Endogenous oxytocin measures are usually collected under basal conditions or during experimental designs in both clinical samples and healthy individuals [2, 6]. An important question that has received relatively little attention is whether oxytocin displays circadian variation similar to the well-established variations of cortisol [7-9]. This question is particularly relevant for methodological aspects and could partially explain significant data variability and mixed findings in the field [6]. However, assessing circadian variation requires at least two or more measurements throughout the day at specific time points, whereas quantification of oxytocin in blood or cerebrospinal fluid may limit this type of assessment in large samples [7, 10, 11]. On the other hand, while the methodological reliability of salivary oxytocin measures is controversial[6], evidence suggests that repeated measurements using radioimmunoassay (RIA) can enhance the reliability of salivary oxytocin assessment [6, 12-14].

Existing evidence regarding oxytocin’s circadian rhythm remains scarce and inconclusive. Van Dam and colleagues [7] assessed salivary oxytocin in 12 adolescents at multiple times during the day and reported a sharp reduction immediately after awakening followed by stable levels during the day. Kagerbauer and colleagues [11] measured oxytocin in cerebrospinal fluid (CSF), blood, and saliva at four time points in 20 neurosurgical patients, and reported circadian variations (decrease between early and late afternoon) only in CSF among postmenopausal women, but not in other participant subgroups or sampling methods. Both studies were well-designed but limited by small sample sizes and heterogeneous participant groups, particularly regarding sex and hormonal status. These aspects are important since menstrual cycle phase and hormonal contraception influence oxytocin secretion due to its interactions with estrogen and progesterone [15]. Graugaard-Jensen et al. [10] addressed some of these limitations by comparing plasma oxytocin levels between naturally cycling women and women using oral contraceptives and found no significant circadian variation. However, morning samples were not collected immediately after individual awakening, as done for cortisol circadian assessment [9].

In the present study we aimed to investigate potential circadian variation in salivary oxytocin levels among young adult women. To address methodological limitations highlighted in previous research, we standardized saliva collection times, recruited a homogeneous sample regarding sex (female only) and age range, assessed all participants during the luteal phase of their menstrual cycle, and accounted for hormonal contraceptive use and antidepressant intake in clinical participants. By implementing these controls, we intended to reduce variability observed in earlier studies and contribute to a clearer understanding of oxytocin’s circadian patterns.

We included both healthy adult women and women diagnosed with borderline personality disorder (BPD). It is of particular interest to study oxytocin in this clinical group because core features of BPD—such as severe emotional dysregulation, interpersonal difficulties, and heightened stress reactivity [16-18], are related to processes modulated by oxytocin, including emotion regulation, stress response, and interpersonal relationships [2, 3, 19]. Prior research on basal oxytocin levels in BPD shows inconsistent results, with some studies reporting lower plasma or serum oxytocin levels and reduced receptor expression compared to healthy controls [20-22], while others found no significant group differences [23, 24]. Additionally, to our knowledge, circadian patterns of oxytocin secretion in BPD have not been explored, with the exception of a pilot investigation conducted by our group [25]. Recently, Bocchio Chiavetto and colleagues [26] reported lower morning plasma oxytocin levels in BPD, although the exact timing of this morning assessment was not specified. Therefore, the present study investigates circadian variation in salivary oxytocin levels in young adult healthy women and women diagnosed with BPD. By employing precise, timed sampling, it aims to further address existing gaps in the literature relevant to both groups

## Methods

The study was approved by the local ethics committee and was registered on ClinicalTrials.gov (NCT05357521). Recruitment and data collection are complete. The protocol included measures of oxytocin and cortisol under naturalistic conditions and during an experimental stress task. The present report focuses on the naturalistic measures of all participants, while and results related to the experimental task will be presented separately since they are related to different research questions.

### Participants

A hundred and twenty-three (123) female participants (61 healthy controls and 62 patients with BPD) were included in the study and signed informed consent. Of these, 91 (47 healthy controls and 44 patients with borderline personality disorder) completed the entire protocol. Healthy controls were recruited by advertisement and patients with BPD within patients treated at the Emotion Regulation Unit of the Division of Psychiatry, Geneva University Hospitals as previously described [27].

Female participants aged 18-35 years were eligible for the study if they were free from neurohormonal or neurological disorders and not using systemic corticosteroids. Exclusion criteria for all participants included pregnancy or breastfeeding within the last 6 months, a history of alcohol or drug addiction, or a body mass index under 17. Participants experiencing major stressful life events (e.g., loss of a close relative, loss of employment, significant changes in marital status or medical condition) within the two months prior to enrolment were excluded initially or contacted for eligibility re-assessment after a few months. Alcohol, tobacco, and occasional cannabis use were permitted, but use was prohibited on sampling days and controlled through interviews, with sampling rescheduled if necessary.

For the BPD group, participants were required to meet DSM-5 criteria, confirmed via structured clinical interview. They were excluded if they had a formal diagnosis of psychosis, were daily users of neuroleptics, or were using antidepressants outside of the SSRI or SNRI classes. Healthy control women were excluded if they met three or more BPD diagnostic criteria (the threshold for a formal diagnosis being five).

### Study Procedures

The entire protocol comprised three visits.

- Visit 1: During their initial visit, participants provided informed consent, underwent eligibility screening, completed a structured clinical interview, and filled out psychometric questionnaires.
- Visit 2: Naturalistic Data Collection: This visit involved the collection of naturalistic data over a single weekday. Participants collected salivary samples at six predetermined time points: upon awakening, 30 minutes post-awakening, 45 minutes post-awakening, between 12:00 and 2pm (early afternoon) in the laboratory, at 6pm, and immediately before sleep. To enhance adherence, participants received a reminder the evening before the sampling day and were instructed to precisely record the time of each collection. For women not using hormonal contraception, Visits 2 and 3 were scheduled to coincide with the luteal phase of their menstrual cycle, ideally one week prior to the expected onset of menstruation. Participants tracked their cycles and reported their average length to allow for individual estimation of the mid-cycle and calculation of the ideal luteal phase timeframe (six days subtracted from the expected end of their cycle). If menstruation began before a scheduled Visit 2 or 3, data collection was rescheduled to maintain cycle consistency. The selection of the luteal phase was based on prior research examining salivary oxytocin and cortisol interactions [28, 29] and the broader aim of facilitating future inclusion of male participants by focusing on a phase with more similar stress responses between sexes compared to the follicular phase [30].
- Visit 3: The third visit was scheduled for the day after Visit 2, or at most four days later. This visit included an experimental stress task (data not reported in this paper) and the return of the evening salivary samples collected during the naturalistic assessment day.

### Hormone Sample Collection and Processing

Saliva samples were collected using Salivettes with a synthetic swab (Sarstedt, Germany), which participants were instructed to chew for at least one minute. Following collection, swabs were temporarily stored at -20°C. For longer-term storage until analysis, samples were centrifuged and stored at -80°C. Sample temperature was maintained with dry ice during shipping.

### Hormonal Measures

Salivary concentrations of oxytocin and cortisol were quantified from the collected samples. Oxytocin: Salivary oxytocin concentrations (expressed in pg/ml) were quantified by radioimmunoassay (RIAgnosis, Sinzing, Germany). Consistent with the planned protocol and considering the lack of well-established validity for long-term oxytocin stability at room temperature or with home refrigeration (relevant for evening samples returned at a later visit), oxytocin was measured only at the first morning time point (upon awakening) and the fourth time point in the early afternoon (collected between 12:00 and 2pm in the laboratory). The saliva samples collected at these two time points were split into two vials each to allow for the measurement of both oxytocin and cortisol from the same biological material. Detailed procedures for oxytocin quantification followed those described in previous studies, including our own [19, 25, 29].

Cortisol: Salivary cortisol concentrations (expressed in Concentration (ug/dl)) were quantified (FCBG, Geneva, Switzerland), at all six collection time points using a direct ELISA kit for human salivary cortisol from IBL Tecan with a mean intra-assay variability of 4.3 % and a mean inter-assay variability of 13.2 %.

### Statistical Analysis

Statistical analyses were conducted using SPSS 28.0. Oxytocin levels were normally distributed at both time-points (Shapiro-Wilk test, p > 0.05 for both time points) and no outliers were identified (SPSS 1.5 interquartile range rule). For oxytocin analyses, a mixed 2 (Group: females with BPD vs. Healthy female controls) × 2 (Time: Awakening, early afternoon) repeated-measures ANOVA was performed to assess circadian variation in salivary oxytocin levels and group differences across time. Effect sizes were reported using partial eta squared (η p2). Box’s test of equality of covariance matrices indicated that the assumption of homogeneity was met (p > 0.05). To explore the potential influence of hormonal contraception, a repeated-measures ANCOVA was conducted, including contraceptive use as a covariate. Given that antidepressant use was exclusive to the BPD group, a separate repeated-measures ANOVA was conducted within this group to test whether antidepressant use influenced circadian changes in salivary oxytocin. All statistical tests were two-tailed, with a significance level set at p < 0.01.

For cortisol analyses, a mixed 2 (Group: females with BPD vs. Healthy female controls) × 6 (Time: Awakening, 30 min post-awakening, 45 min post-awakening, early afternoon, 6pm and Bedtime) repeated-measures ANOVA was performed.

## Results

### Sample characteristics

As shown in Tables 1 and 2, groups did not significantly differ regarding age, body mass index (BMI), use of hormonal contraception, or cycle length

**Table 1.** Group characteristics.

|  | **Females with BPD**<br>**(n = 44)** | **Healthy female**<br>**controls (n = 47)** | **Test Statistic** | ***P***<br>**value** |
| --- | --- | --- | --- | --- |
| Age (years) | 24.4 ± 5.07 (18–35) | 25.1 ± 4.38 (18–35) | U = 767<br>(Mann–Whitney) | 0.46 |
| Body Mass<br>Index | 24.0 ± 5.35 (17.5–46.9) | 22.2 ± 3.74 (18.0–<br>41.7) | U = 646<br>(Mann–Whitney) | 0.06 |
| Hormonal<br>contraception | 12 (27.3%) | 15 (31.9%) | $\chi^2(1) = 0.49$ | 0.63 |

*BPD* Borderline Personality Disorder. M ± SD (range).
*BPD* Borderline Personality Disorder.

**Table 2.** Cycle Duration in natural Cycling women.

| **Cycle Duration**<br>**(days)** | **Females with BPD**<br>**(n = 32)** | **Healthy female controls**<br>**(n = 32)** |
| --- | --- | --- |
| < 26 | 3 | 4 |
| 27–30 | 18 | 19 |
| 30–35 | 9 | 8 |
| > 35 | 2 | 1 |

### Oxytocin

Figure 1 shows salivary oxytocin levels (pg/ml) at awakening and early afternoon for females with BPD and healthy female controls. The repeated-measures ANOVA revealed a significant main effect of Time (*F* (1,83) = 28.03, *p* < 0.001, η^2^p = 0.25), indicating that salivary oxytocin levels significantly differed between awakening and early afternoon. There was no significant main effect of Group (*F* (1, 83) = 3.23, *p* =0.08), nor significant Group × Time interaction (*F* (1,83) = 1.17, *p* = 0.28). As salivary oxytocin was missing at one of the two time-points for one healthy female and five females with BPD (due to low amounts of saliva after centrifugation), we imputed the missing values with the corresponding available time-point value (a conservative approach) and ran the ANOVA with these imputed values. Effects of time remained significant (*F* (1, 89) = 26.80, *p* <0.001), with no group effects or Group X time interaction.

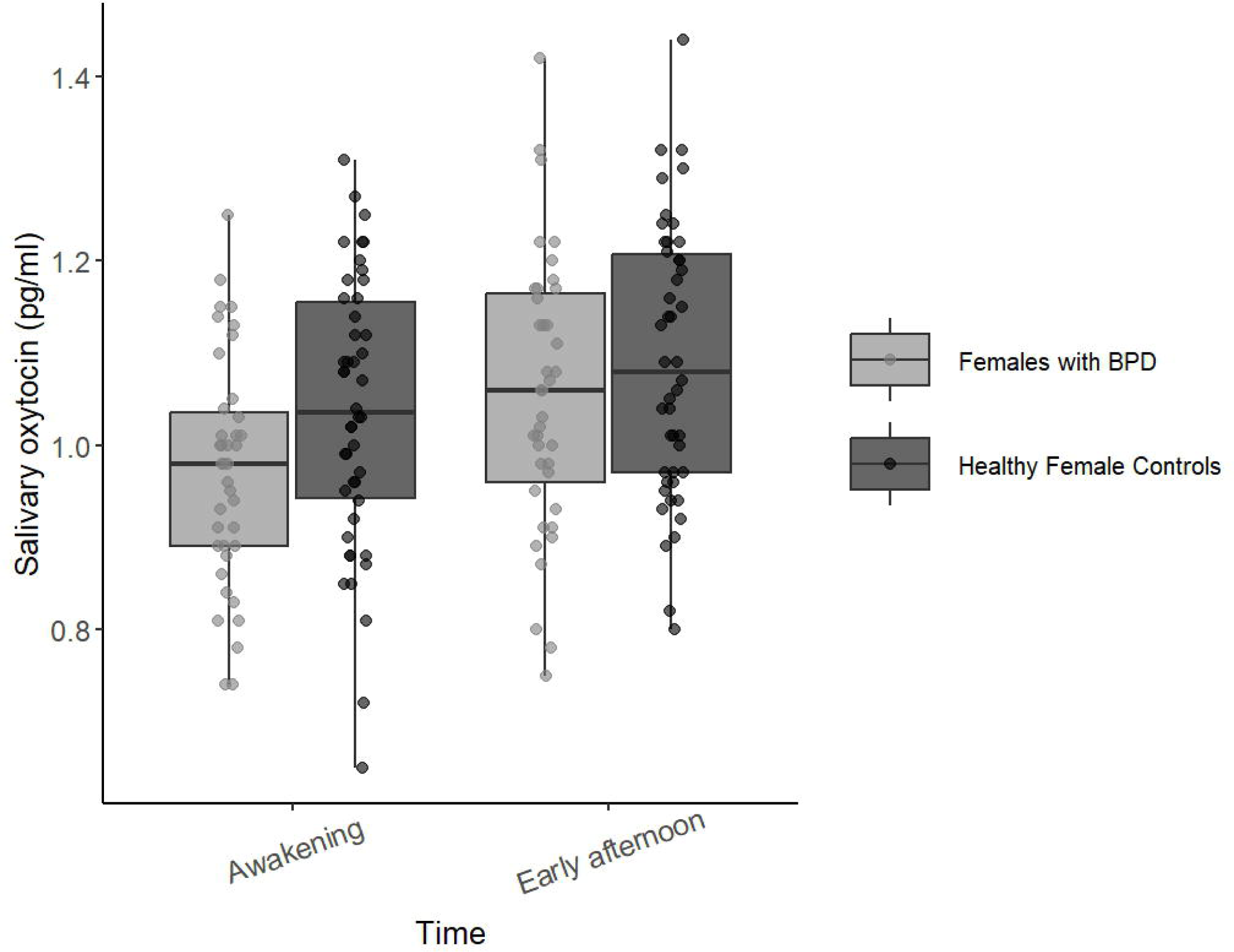

A total of 27 women in the entire sample (15 healthy female controls and 12 patients with BPD) reported the use of hormonal contraception (see Table 3 for a detailed description of contraception type). When hormonal contraception was included as a covariate, the main effect of Time remained significant (*F* (1,82) = 14.5, *p* < 0.01), with no significant main effect of group or Group X Time interaction.

**Table 3.**
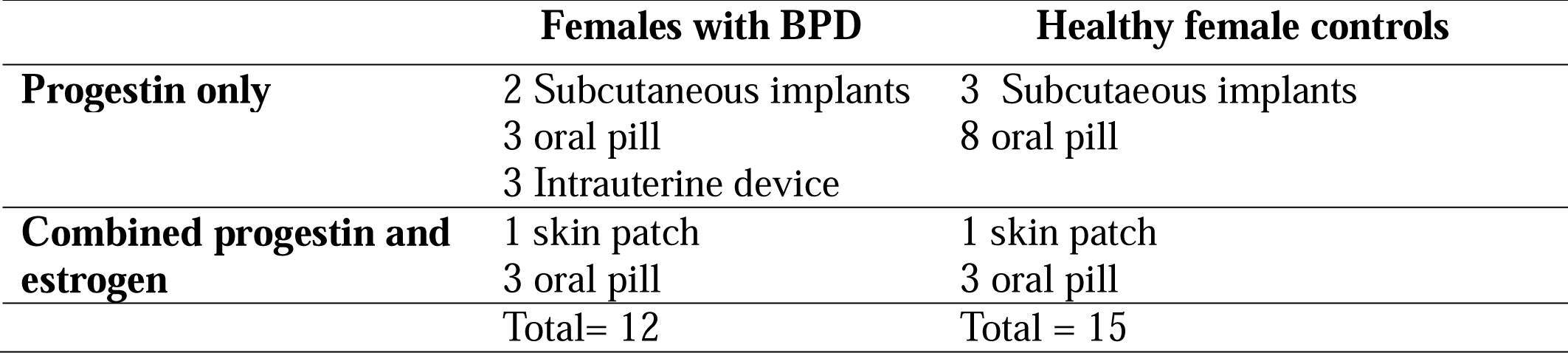
Hormonal contraception: type and administration method.

|  | **Females with BPD** | **Healthy female controls** |
| --- | --- | --- |
| **Progestin only** | 2 Subcutaneous implants<br>3 oral pill<br>3 Intrauterine device | 3 Subcutaneous implants<br>8 oral pill |
| **Combined progestin and estrogen** | 1 skin patch<br>3 oral pill | 1 skin patch<br>3 oral pill |
|  | Total= 12 | Total = 15 |

Among women with BDP, 11 were treated with an SSRI antidepressant. A repeated-measures ANOVA showed a significant main effect of time (*F* (1, 37) = 17.84, *p* < 0.001), but no main effect of group (*F* (1, 37) = 0.85, *p* = 0.36) or Time X Antidepressant interaction (*F* (1, 37) = 1.15, *p* = 0.29).

### Cortisol

Figure 2 shows salivary cortisol levels across time points for females with BPD and healthy female controls. The repeated-measures ANOVA revealed a significant main effect of Time (*F* (5,78) = 44.21, *p* < 0.001, η^2^p =0.36), indicating that salivary cortisol levels significantly varied across time. There was no significant main effect of Group (*F* (1, 78) = 2.09, *p* = 0.15), nor significant Group × Time interaction (*F* (5, 78) = 0.25, *p* = 0.96). When missing values were imputed (eight at awakening, two at mid-day and 1 at 6pm), the effect of time remained significant, with no main effect of group or interaction.

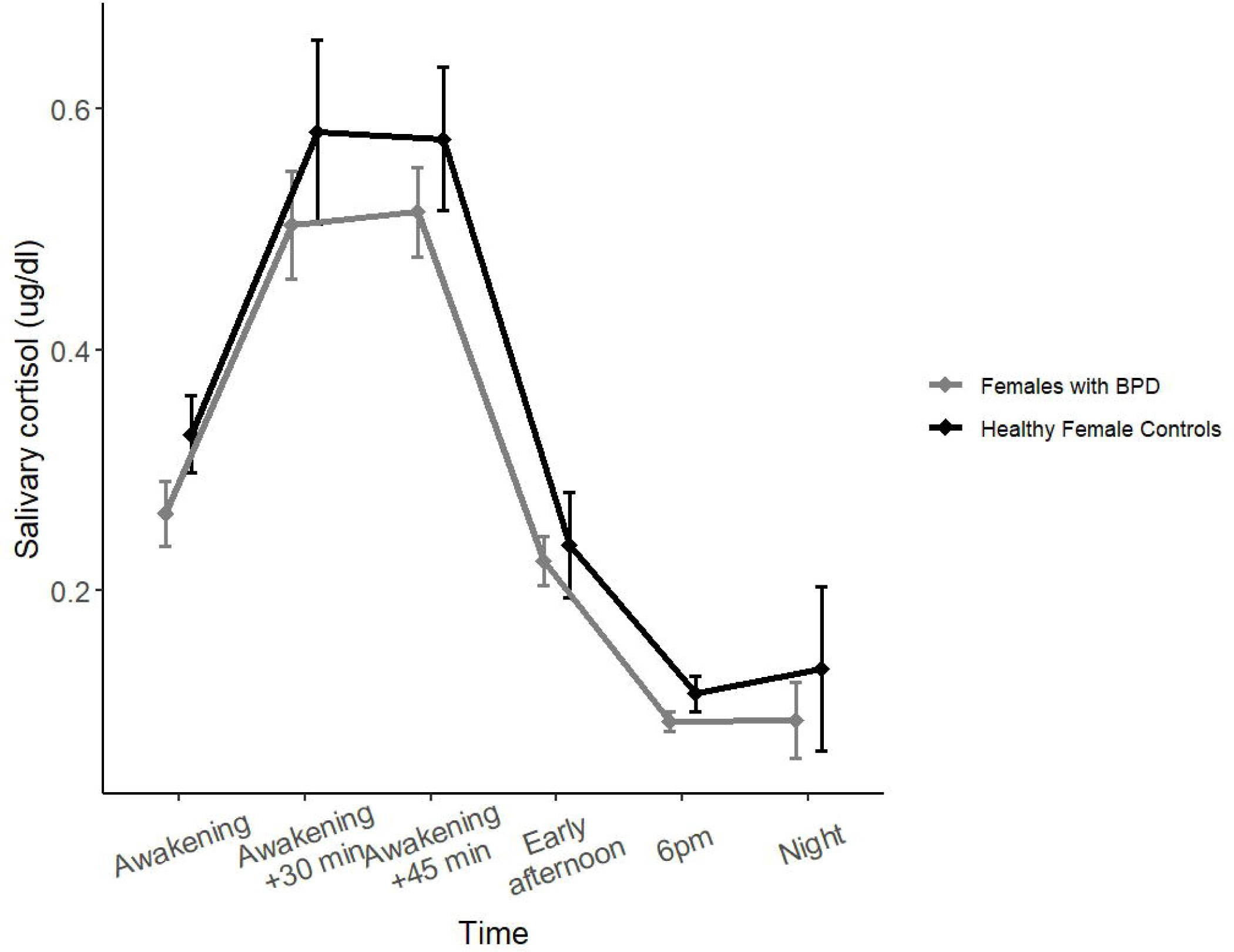

## Discussion

This study investigated potential circadian variation in salivary oxytocin levels among young adult women, including patients with Borderline Personality Disorder (BPD) and healthy controls. Across the entire sample, we observed a significant increase in salivary oxytocin from awakening to early afternoon. While there were no significant group differences or interactions, we observed a non-significant trend (*p* = 0.08) toward lower oxytocin levels in the BPD group. Overall findings indicate the importance of considering the time of day in studies utilizing salivary oxytocin as a measure. The subsequent discussion will interpret these results within the context of existing literature and explore their implications for understanding oxytocin dynamics.

Previous studies investigating the diurnal variation of oxytocin levels showed inconsistent findings. Some studies indicated circadian patterns, such as a decline at awakening with stable daytime levels [7] or changes observed from morning to afternoon in specific populations (e.g., postmenopausal women) [11]. In contrast, absence of significant diurnal oxytocin variation was also reported [10]. The differences between our findings and those of prior studies may be related to methodological variations. Our study included a larger and relatively homogeneous sample of young adult women and controlled for factors such as menstrual cycle phase, hormonal contraception, and medication use. Furthermore, the specific time points for sample collection in our protocol (awakening and early afternoon) differed from some previous studies. Another distinction is that our study utilized a non-hospitalized sample assessed within participants’ natural environment, whereas some previous studies employed in-patient protocols [11]. In contrast, absence of significant diurnal oxytocin variation was also reported [10]. The differences between our findings and those of prior studies may be related to methodological variations. Our study included a larger and relatively homogeneous sample of young adult women and controlled for factors such as menstrual cycle phase, hormonal contraception, and medication use. Furthermore, the specific time points for sample collection in our protocol (awakening and early afternoon) differed from some previous studies. Another distinction is that our study utilized a non-hospitalized sample assessed within participants’ natural environment, whereas some previous studies employed in-patient protocols [7, 10, 11]. This approach may have helped to minimize potential confounds associated with the stress of a hospital setting or concurrent medical interventions, factors that can influence oxytocin release [19, 28, 29, 31].

Beyond the observed circadian pattern, we noted a non-significant trend towards lower salivary oxytocin levels in the Borderline Personality Disorder group compared to healthy controls (p = 0.08). This observation presents a point of interest when considered alongside previous studies that have reported lower plasma oxytocin levels under basal conditions in individuals with BPD [20-22, 26]. Direct comparisons are challenging due to differences in the type of measurement (saliva vs. plasma or receptor expression) and potential variations in the clinical characteristics of the BPD samples, as well as methodological factors in sample collection and analysis. While our finding did not reach statistical significance, it aligns with the direction of effects observed in some previous plasma-based studies, suggesting this warrants further investigation in larger salivary oxytocin studies in this population.

Within the BPD group, a subset of participants were treated with antidepressant medication, restricted to Selective Serotonin Reuptake Inhibitors (SSRIs). Subgroup analyses comparing BPD participants on SSRIs versus those not on SSRIs did not reveal significant differences in salivary oxytocin circadian variation or overall levels. However, it is important to acknowledge the inherent heterogeneity within the BPD diagnosis, which encompasses a wide range of clinical profiles, symptom severity, and comorbidity. Given this variability, a more detailed characterization of clinical profiles in future studies could provide valuable insights into potential associations with oxytocin dynamics. Furthermore, exploring the effects of other classes of psychotropic medications (not included in our current study) would merit investigation.

In addition to examining group differences, we accounted for potential confounding variables known to influence oxytocin levels [1, 15]. The groups did not significantly differ with respect to age, menstrual cycle phase, or the use of hormonal contraception. While hormonal contraception was included as a covariate in our analyses, we did not observe a significant effect of hormonal contraception on the circadian variation of salivary oxytocin, nor a significant interaction with group. However, it is important to note that previous research suggests that hormonal contraception can influence oxytocin release [15, 32]. Given these findings and the potential for hormonal fluctuations to impact oxytocin dynamics, continued assessment and control for variables such as age, menstrual cycle phase, and hormonal contraception use remain important considerations in future studies investigating oxytocin.

This study presents certain limitations that warrant consideration. Foremost, the sample was restricted to young adult women aged 18 to 35 years during the luteal phase of the menstrual cycle. Consequently, our findings may not be generalizable to males, women in other menstrual cycle phases, or premenarchal or postmenopausal individuals. While the luteal phase was selected for data collection to facilitate comparison with previous research examining oxytocin and cortisol interactions [28, 29], we recognize that this phase is characterized by higher fluctuations in sex hormones, which may introduce variability [15, 33]. This contrasts with some studies that have focused on assessing oxytocin during the more stable follicular phase [e.g., Jobst et al.,]. Furthermore, our estimation of menstrual cycle phase was based on self-report, and we lacked direct hormonal measures of estrogen and progesterone. Given the known interactions between sex hormones and oxytocin [15, 33], future studies including objective hormonal profiling would provide a more accurate understanding. Additionally, although the current sample size was sufficient to detect within-subject circadian variation, future protocols designed to comprehensively assess circadian oxytocin dynamics could benefit from including a greater number of sampling time points throughout the day, potentially extending to later assessments, similar to standard cortisol protocols. However, this approach faces challenges related to the stability of salivary oxytocin without appropriate refrigeration, particularly for evening samples in out-patient contexts.

Despite these limitations, our findings yield important methodological and theoretical implications. The presence of circadian variation in salivary oxytocin levels in young adult women underscores the necessity for future studies to carefully control for the time of day at which samples are collected. Implementing such standardized collection protocols can reduce methodological variability and thereby enhance the replicability and interpretability of oxytocin research findings. Conversely, a failure to account for these diurnal fluctuations may contribute to inconsistencies observed across studies

## References

1. Audunsdottir, K. and D.S. Quintana, Oxytocin’s dynamic role across the lifespan. Aging Brain, 2022. 2: p. 100028.

2. Carter, C.S., et al., Oxytocin: Not “just a female hormone”: A very special issue and a very special molecule. Compr Psychoneuroendocrinol, 2024. 20: p. 100273.

3. Quintana, D.S. and A.J. Guastella, An Allostatic Theory of Oxytocin. Trends Cogn Sci, 2020. 24(7): p. 515–528.

4. Yao, S. and K.M. Kendrick, How does oxytocin modulate human behavior? Mol Psychiatry, 2025.

5. Kirsch, P., Oxytocin in the socioemotional brain: implications for psychiatric disorders. Dialogues Clin Neurosci, 2015. 17(4): p. 463–76.

6. Martins, D., et al., Salivary and plasmatic oxytocin are not reliable trait markers of the physiology of the oxytocin system in humans. Elife, 2020. 9.

7. Van Dam, J.M., et al., Variability of the cortisol awakening response and morning salivary oxytocin in late adolescence. J Neuroendocrinol, 2018. 30(11): p. e12645.

8. Ryan, R., et al., Use of Salivary Diurnal Cortisol as an Outcome Measure in Randomised Controlled Trials: a Systematic Review. Ann Behav Med, 2016. 50(2): p. 210–36.

9. Stalder, T., et al., Evaluation and update of the expert consensus guidelines for the assessment of the cortisol awakening response (CAR). Psychoneuroendocrinology, 2022. 146: p. 105946.

10. Graugaard-Jensen, C., et al., Oral Contraceptives and Renal Water Handling: A diurnal study in young women. Physiol Rep, 2017. 5(23).

11. Kagerbauer, S.M., et al., Absence of a diurnal rhythm of oxytocin and arginine-vasopressin in human cerebrospinal fluid, blood and saliva. Neuropeptides, 2019. 78: p. 101977.

12. Leng, G. and N. Sabatier, Measuring Oxytocin and Vasopressin: Bioassays, Immunoassays and Random Numbers. J Neuroendocrinol, 2016. 28(10).

13. Martin, J., et al., Oxytocin levels in saliva correlate better than plasma levels with concentrations in the cerebrospinal fluid of patients in neurocritical care. J Neuroendocrinol, 2018: p. e12596.

14. Tabak, B.A., et al., Advances in human oxytocin measurement: challenges and proposed solutions. Mol Psychiatry, 2023. 28(1): p. 127–140.

15. Quintana, D.S., et al., The interplay of oxytocin and sex hormones. Neurosci Biobehav Rev, 2024. 163: p. 105765.

16. Association, A.P., Diagnostic and statistical manual of mental disorders (5th ed.).. 2013: Arlington.

17. Meehan, K.B., J.F. Clarkin, and M.F. Lenzenweger, Conceptual Models of Borderline Personality Disorder, Part 1: Overview of Prevailing and Emergent Models. Psychiatr Clin North Am, 2018. 41(4): p. 535–548.

18. Paris, J., Differential Diagnosis of Borderline Personality Disorder. Psychiatr Clin North Am, 2018. 41(4): p. 575–582.

19. Jong, T.R., et al., Salivary oxytocin concentrations in response to running, sexual self-stimulation, breastfeeding and the TSST: The Regensburg Oxytocin Challenge (ROC) study. Psychoneuroendocrinology, 2015. 62: p. 381–8.

20. Bertsch, K., et al., Reduced plasma oxytocin levels in female patients with borderline personality disorder. Horm Behav, 2013. 63(3): p. 424–9.

21. Carrasco, J.L., et al., Decreased oxytocin plasma levels and oxytocin receptor expression in borderline personality disorder. Acta Psychiatr Scand, 2020. 142(4): p. 319–325.

22. Ebert, A., et al., Endogenous oxytocin is associated with the experience of compassion and recalled upbringing in Borderline Personality Disorder. Depress Anxiety, 2018. 35(1): p. 50–57.

23. Bomann, A.C., et al., The neurobiology of social deficits in female patients with borderline personality disorder: The importance of oxytocin. Personal Ment Health, 2017. 11(2): p. 91–100.

24. Bonfig, J., S.C. Herpertz, and I. Schneider, Altered hormonal patterns in borderline personality disorder mother-child interactions. Psychoneuroendocrinology, 2022. 143: p. 105822.

25. Aboulafia-Brakha, T., et al., Hypomodulation of salivary oxytocin in patients with borderline personality disorder: A naturalistic and experimental pilot study. Psychiatry Research Communications, 2023. 3(2).

26. Bocchio Chiavetto, L., et al., Reduction of oxytocin plasma levels in borderline personality disorder and normalization induced by psychotherapies. Psychol Med, 2025. 55: p. e92.

27. Blay, M., et al., Body modifications in borderline personality disorder patients: prevalence rates, link with non-suicidal self-injury, and related psychopathology. Borderline Personal Disord Emot Dysregul, 2023. 10(1): p. 7.

28. Alley, J., et al., Associations between oxytocin and cortisol reactivity and recovery in response to psychological stress and sexual arousal. Psychoneuroendocrinology, 2019. 106: p. 47–56.

29. Bernhard, A., et al., Adolescent oxytocin response to stress and its behavioral and endocrine correlates. Horm Behav, 2018. 105: p. 157–165.

30. Kirschbaum, C., et al., Impact of gender, menstrual cycle phase, and oral contraceptives on the activity of the hypothalamus-pituitary-adrenal axis. Psychosom Med, 1999. 61(2): p. 154–62.

31. Engert, V., et al., Boosting recovery rather than buffering reactivity: Higher stress-induced oxytocin secretion is associated with increased cortisol reactivity and faster vagal recovery after acute psychosocial stress. Psychoneuroendocrinology, 2016. 74: p. 111–120.

32. Garforth, B., et al., Elevated plasma oxytocin levels and higher satisfaction with life in young oral contraceptive users. Sci Rep, 2020. 10(1): p. 8208.

33. Schmalenberger, K.M., et al., How to study the menstrual cycle: Practical tools and recommendations. Psychoneuroendocrinology, 2021. 123: p. 104895.

